# SNP-based quantitative deconvolution of biological mixtures: application to the detection of cows with subclinical mastitis by whole genome sequencing of tank milk

**DOI:** 10.1101/740894

**Authors:** Wouter Coppieters, Latifa Karim, Michel Georges

**Affiliations:** Genomics Platform, GIGA Institute, University of Liège; Unit of Animal Genomics, GIGA Institute & Faculty of Veterinary Medicine, University of Liège

## Abstract

Biological products of importance in food (f.i. milk) and medical (f.i. donor blood derived products) sciences often correspond to mixtures of samples contributed by multiple individuals. Identifying which individuals contributed to the mixture and in what proportions may be of interest in several circumstances. We herein present a method that allows to do this by shallow whole genome sequencing of the DNA in mixed samples from hundreds of donors. We demonstrate the efficacy of the approach for the detection of cows with subclinical mastitis by analysis of farms’ tank mixtures containing milk from as many as 500 cows.

## Introduction

Mastitis, <u>i.e. the inflammation of the udder,</u> is the most important health issue in dairy cattle<u>. It is estimated to</u> cost European farmers > 1 billion € per year in treatment and milk loss <u>(Hogeveen et al., 2011). Upon inflammation, immune cells migrate in the udder and milk. While milk from healthy cows typically contains > 100,000 cells per milliliter (ml) of milk, these numbers (referred to as Somatic Cell Counts or SCC) typically increase into the millions in case of mastitis. Prior to the manifestation of overt clinical symptoms, SCC progressively increase in the milk of cows developing mastitis: SCC</u> ≥ <u>200,000 are typically considered to be a sign of pre-or sub-clinical mastitis. Both yield and quality of the milk of cows with subclinical mastitis is reduced (Schukken et al., 2003). M</u>astitis is routinely managed by periodically counting <u>SCC</u> in milk samples to preemptively identify cows developing subclinical udder inflammation. As profit margins decrease, farmers tend to forgo milk testing thereby compromising health management. Cost-effective alternatives for rapid detection of cows with subclinical mastitis are <u>hence</u> neede<u>d (Viguier et al., 2009)</u>.

<u>The milk obtained from individual cows is typically collected in one or more large “milk tanks” on the farm, before being shipped to dairy factories.</u> We previously proposed that somatic cell counts (SCC) in the milk of individual cows could be estimated <u>by measuring</u> the allel<u>ic</u> frequencies <u>in the tank milk</u> for sufficient numbers of SNPs, provided that all cows contributing milk to the tank be genotyped <u>once</u> for the corresponding variants. Thus, the proposed method would allow <u>the identification of</u> a minority of cows with subclinical mastitis by <u>regularly</u> analyzing a single sample containing a mixture of milk from all the cows on the farm, hence dramatically reducing costs. <u>Prior to ~2010 estimation of breeding values to select the best dairy sires and dams used pedigree-based estimates of kinship. Since then, selection methods increasingly use genome-wide SNP information in a process referred to as “genomic selection” (GS) (Georges et al., 2019).</u> As GS is becoming routine <u>in dairy cattle</u> (including for dams), herds that are fully genotyped with <u>genome-wide SNP</u> arrays are becoming standard, and the proposed method feasible. We herein demonstrate that by combining low density SNP genotyping or shallow sequencing of the cows and tank milk’s DNA with in silico genotype imputation, individual SCC can be accurately determined and cows with subclinical mastitis effectively identified even in the largest farms (≥ 500). The proposed method has the potential to dramatically improve the monitoring of udder health in dairy farms, and to allow the tracing of the origin of bulk animal food products other than milk.

## Results

### Principle of the proposed method

Assume that cows and tank (i.e. the reservoir in which the milk of the cows is collected) milk are genotyped for a collection of SNPs. <u>Assume that the interrogated SNPs are biallelic, each characterized by a *A* (say the allele of the reference genome) and a *B* allele (say the alternate allele).</u> If all cows contribute identical amounts of DNA to the milk, the expected <u>proportion of the *B* allele (commonly referred to as “B-allele frequency” when analyzing SNP array data particularly to search for Copy Number Variants)</u> in the tank milk corresponds to the frequency of the *B* allele in the farm’s cow population. The actual DNA amount contributed by each cow depends on the volume of milk <u>that she</u> produced and its SCC. Unequal DNA contributions will cause slight departures from the expected *B* allele frequencies in the tank milk. Integrating these shifts over a large number of SNPs in conjunction with the known genotypes of individual cows allows for the estimation of the relative DNA contribution of each cow. <u>This can for instance be achieved using a set of *m* linear equations in which the “B-allele frequency” of each SNP *j* (of *m*) is modelled as the sum (over *n* cows) of the products of the dosage of the B allele in the genotype of cow *j* (*d_ij_*, known from her SNP genotype) multiplied by the proportion of DNA contributed by cow *i* (*f_i_*) to the milk. The proportions of DNA contributed by each cow can then be estimated using for instance least square methods</u>. Accounting for individual milk volumes and for the SCC in the tank milk allows for the estimation of SCC for individual cows (Fig. 1 and Methods).

**Figure 1:**
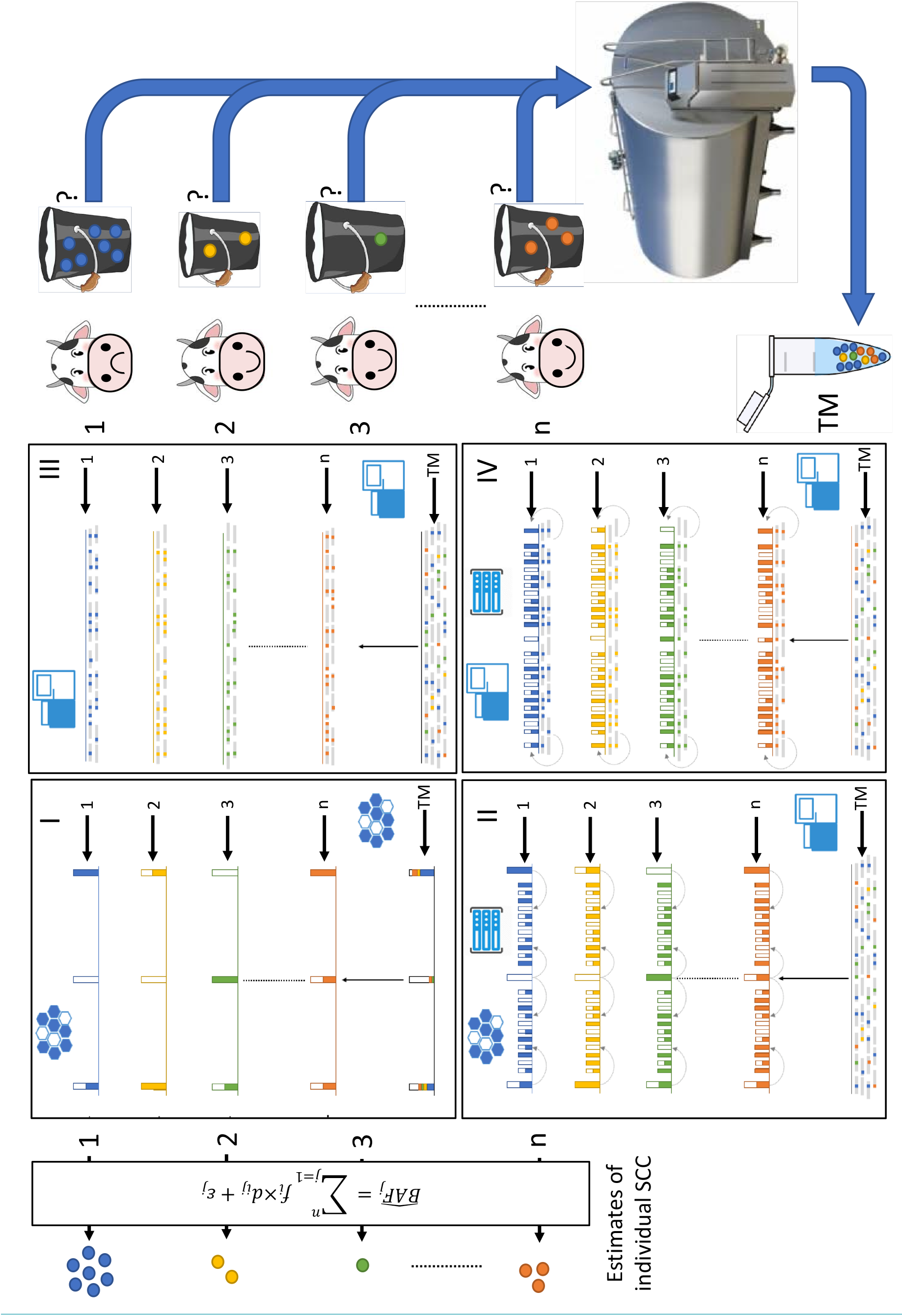
Estimating Somatic Cell Counts (SCC) in the milk of individual cows by analyzing a sample of milk from the farm’s tank. Cows 1 to *n* contribute different amounts of milk (buckets of various sizes in the figure) to the farm’s tank. The milk contains somatic cells (shown as small spheres in the milk colored by cow) whose numbers reflect the health status of the cow’s udder. Cow 1 has higher SCC, an indicator of subclinical mastitis. SCC are unknown upon milking (indicated by the “?”). Cows are individually SNP genotyped once. In scheme <u>I</u> this is done using SNP arrays (illustrated by the mesh) yielding genotype information for the limited number of interrogated SNPs (high bars) that can be summarized by the B-allele frequency as shown (white: 0, halve colored: 0.5, full<u>y</u> colored: 1). SNP genotypes of individual cows are coded in the same colors as the SCC. In scheme <u>II</u>, the genotypes of the interrogated SNPs are augmented by imputation (illustrated by the computer rack), yielding dosage information (B-allele frequency) for many more SNPs (small bars). In scheme <u>III</u>, cows are genotyped individually by shallow whole genome sequencing (SWGS) (illustrated by the sequencer). Sequence reads (gray lines) are aligned to the reference genome and alternate alleles at SNP positions highlighted as color-coded tics. The *B*-allele frequency at specific SNP positions is measured as the ratio of the number of reads with <u>the *B* allele</u> vs the total number of reads. In scheme <u>IV</u>, the genotype information from SWGS is augmented by imputation improving the accuracy of the B-allele frequency estimates for millions of SNPs (small bars). A small sample of milk (T(ank) M(ilk)) is periodically (f.i. monthly or weekly) collected from the farm’s tank. DNA is extracted from TM and genotyped using SNP arrays (scheme <u>I</u>) or SWGS (schemes <u>I</u>, <u>II</u> and <u>IV</u>). B-allele frequency for SNP *j* in the milk 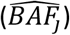 is estimated from the ratio of fluorescence intensities when using SNP arrays, or from the proportion of reads with *B* allele in SWGS. The SCC of individual cows are estimated from a set of linear equations modelling 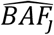 as the sum of *B* allele dosage (*d_ij_*) multiplied by the proportion of the DNA in the tank contributed by cow *i* (*f_i_*). The estimated proportions of DNA contributed by each cow correspond to the values of *f_i_*’s that minimize the sum of squared errors (*ε*) over all SNPs. The SCC for individual cows, *per se*, can be estimated as *SCC_i_* = *SCC_tank_* × *V_tank_* × *f_i_/V_i_*, where *SCC_tank_* is the SCC measured in the farm’s tank, and *V_i_*/*V_tank_* is the proportion of the milk volume contributed by cow *i*

### Evaluating the proposed method by simulation

We first evaluated the proposed method by simulation (cfr. Methods). Genotyping the cows and the tank milk using 10K SNP arrays (i.e. low-density (LD) arrays as generally used <u>in the context of genomic selection</u>) allowed for the accurate estimation of individual SCC for farms with up to 100 cows (*r* ≥ 0.9, where *r* is the correlation between real and estimated SCC) (scheme <u>I</u>). However, farms with > 100 cows are increasingly common. Medium- (MD, f.i. 50K) and high-density (HD, f.i. 700K) SNP arrays would be needed for the approach to be effective in farms with ≥ 250 or ≥ 500 cows, respectively. Yet – being too expensive - this is presently not a viable proposition (Fig. 2A <u>and Supplemental Table 1</u>). We therefore envisaged a second scheme (<u>II</u>) in which the cows would still be genotyped with LD SNP arrays (as done in practice) yet imputed (<u>Marchini & Howie, 2010</u>) to whole genome (8 million SNPs in the simulations) using a sequenced reference population (<u>Daetwyler et al., 2014</u>), while the DNA of the tank milk would by genotyped by shallow whole-genome sequencing (SWGS). We found that under this scenario sequencing the tank milk at a depth of 0.25 was sufficient for farms with 100 cows, 0.5 for farms with 250 cows, and 2 for farms with 500 cows (Fig. 2<u>B</u>). Accuracies were not significantly affected by the density of the SNP arrays, i.e. the method performed as well with LD as with MD arrays (data not shown). Anticipating further advances in sequencing technology, we also envisaged a scheme (<u>III</u>) in which both cows and tank milk would be genotyped by SWGS. We found that a 1-fold sequencing depth of the tank milk would be sufficient when combined with a 0.25-fold depth for 100 cows, while a 5-fold sequencing depth of the tank milk would be needed in combination with 0.25-fold depth for 250 cows and 1-fold depth for 500 cows (Fig. 2<u>C&D</u>). In scheme <u>III</u>, allelic dosage in the cows is directly measured from the number of alternative and reference alleles in the sequence reads. We further explored the effectiveness of augmenting the cow genotype information from SWGS by imputation (scheme <u>IV</u>). This proved to be effective, reducing the required sequence depth to 0.25-fold for tank milk and 0.25-fold for 100 cows, to 1-fold for tank milk and 0.25-fold for 250 cows, and to 5-fold for tank milk and 0.25-fold for 500 cows (Fig. 2). <u>The previous simulations make a number of assumptions that may not apply in the real world: (i) SNPs were sampled from a uniform distribution (i.e. rare and common SNPs equally represented), (ii) SNPs were assumed to be in linkage equilibrium, (iii) cows on the farm were assumed to be unrelated, and (iv) milk volumes were assumed to be known without error. To more accurately mimic real conditions we repeated the simulations by (i) sampling genotypes from a phased dataset of 750 Holstein-Friesian whole genome sequences (hence properly accounting for true MAF distribution, true linkage disequilibrium (LD) and relatedness - many of the sequenced animals were related as parent offspring, full- or half-sibs), and (ii) adding a normally distributed error with mean 0 and standard deviation of five liter to the simulated milk volumes (normally distributed with mean of 30 liter and standard deviation of 10 liter). This error rate corresponds approximately to that expected when having to estimate the daily milk volume from the total lactation yield using a standard lactation curve (Miel Hostens, personal communication). We assumed in these simulations that the genotypes of the cows were known without error and that the milk was sequenced at a depth ranging from 0.25 to 5 as before. MAF, LD and relatedness jointly had a relatively modest impact on the accuracy of the method, which could be compensated for by increasing the sequencing depth of the milk to five-fold and still allowing for accurate estimates even in farms with 500 cows. Estimating the milk volume with error had a more pronounced impact on the accuracy making it possible difficult to reach a correlation reaching 0.9 in farms with 500 cows (Fig. 2)</u>.

**Figure 2:**
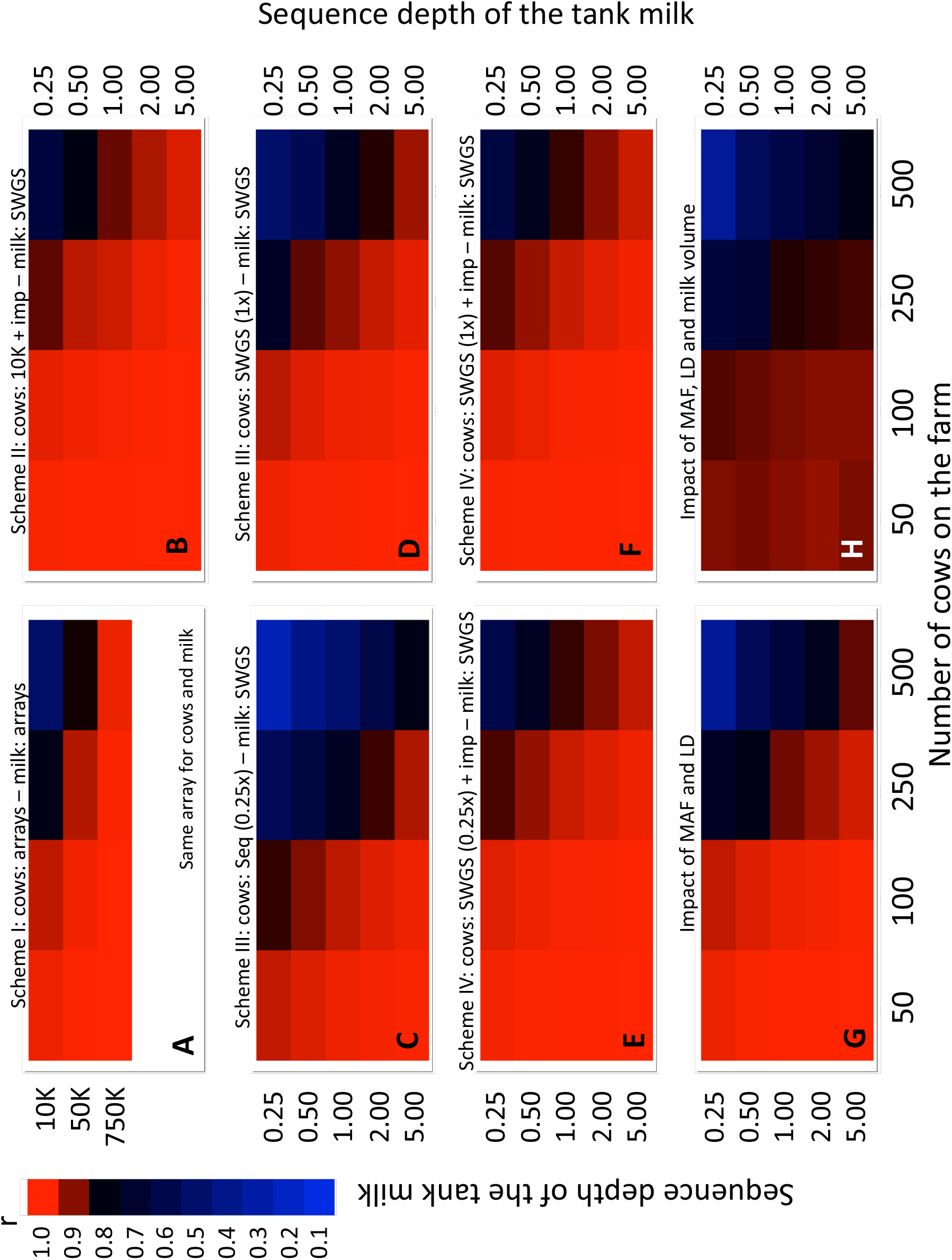
Evaluating the efficiency of the proposed approach by simulation. **(A)** Reference scheme ļ_in which individual cows and tank milk are genotyped with the same array interrogating 10K (LD), 50K (MD) or 700F (HD) SNPs. **(B)** Scheme <u>II</u> in which individual cows are genotyped with a LD 10K SNP array and imputed to whole-genome (8 million SNPs), while the tank milk is whole-genome sequenced at depth ranging from 0.25x to 5x. **(C)** Scheme <u>III</u> which individual cows (<u>0.25×</u>) and tank milk (range: 0.25x to 5x) are genotyped by shallow whole-genome sequencing (SWGS). <u>(D) Same as C except that individual cows are sequenced at 1x depth. (E)</u> Scheme <u>IV</u> in which individual cows are genotyped by SWGS (0.25x) followed by imputation to whole genome (8M SNPs), and tank milk is genotyped by SWGS (range: 0.25x to 5x). <u>(F) Same as E except that individual cows are sequenced at 1x depth. (G) Scheme in which the cow genotypes are sampled from a real dataset hence conform to reality with regards to distribution of MAF, LD and relatedness. Genotypes of the cows are assumed to be known (very similar to II and IV) and tank milk genotyped by SWGS (range: 0.25x to 5x). (H) Same as G except that the milk volume is estimated with error. The color code used to quantify the correlations between predicted and real SCC is shown. Corresponding numerical values are provided in Suppl. Table 1</u>

### Real-world application of the proposed method

To test the feasibility of our method in the real world, we first collected cow (blood) and tank (milk) samples from a farm milking 133 Holstein-Friesian cows. When only using genotypes from the Illumina LD arrays (17K SNPs) for both cows and tank milk (scheme A), correlations between predicted and measured SCC were 0.91 (or 0.79 when ignoring one cow with SCC > 3 million). We then imputed the cows to whole genome (13M SNPs) using a reference population of ~750 whole genome sequenced Holstein-Friesian animals, and sequenced the tank milk at ~3.5-fold depth. The corresponding correlations (scheme B) were 0.97 (0.95) when using all sequence information, or 0.96 (0.92) when down-sampling sequence information as low as 0.1-fold depth (Fig. 3A). We next performed a similar experiment on a farm milking 520 Holstein-Friesian cows. The correlation between predicted and measured SCC was 0.78 (or 0.42 when ignoring 23 cows with SCC > 3 million) when only using information from the LD array for both cows and tank milk (scheme A). When imputing the cows to whole genome (13M SNPs) and sequencing the milk at ~3.5-fold depth (scheme B), the correlation increased to 0.89 (0.83). Down-sampling the sequence information to 0.1-fold depth reduced the correlation to 0.79 (0.57) (Fig. 3B).

**Figure 3:**
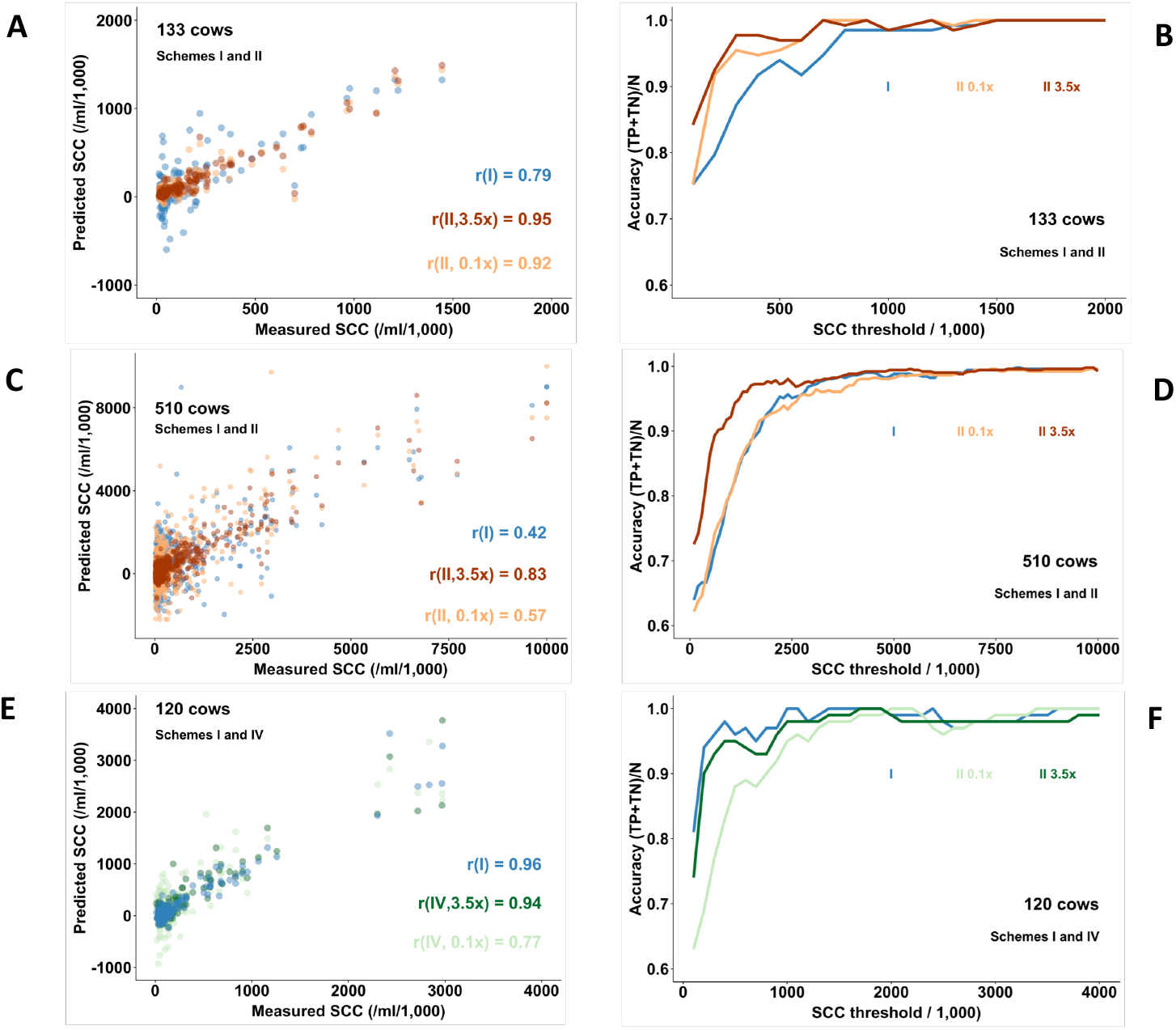
Correlation between predicted and measured SCC in the milk of individual cows (<u>A,C,E</u>), as well as accuracies in classifying cows with SCC above and below a chosen threshold value (<u>B,D,F</u>), in farms with 133 (<u>A,B</u>), 520 (C,D) and 120 (<u>E,F</u>) cows, using scheme <u>I</u> (blue), scheme <u>II</u> (red), or scheme <u>IV</u> (green). Scheme <u>I</u>: cows and tank milk genotyped with LD SNP arrays (17K), no imputation. Scheme <u>II</u>: cows genotyped with LD array and imputed to 13M SNPs, tank milk sequenced 3.5x (red) or 0.1x (orange). Scheme <u>IV</u>: cows genotyped by whole-genome sequencing (1x) and imputation to HD, and tank milk sequenced at 3.5x (dark green) or 0.1x (light green).

As shown in both farms, correlation estimates are affected by SCC spread: small numbers of cows with very high SCC tend to inflate *r*. We therefore computed accuracies, computed as the proportion of correctly classified cows for different SCC thresholds, which is how farmers would likely use the information. It can be seen that for a threshold value of for example 500,000 SCC, accuracies > 0.85 were obtained when sequencing (scheme B) the tank milk at respectively 0.1x (133 cows) and 3.5x depth (520 cows). Thus - as predicted by the simulations - scheme A provided adequate precision for the farm with 133 cows, but not for the farm with 520 cows. However, in this large farm, combining SWGS of the tank milk with whole genome imputation of the cows (i.e. scheme B) was indeed effective (Fig. 3).

As costs per bp continue to decline, sequencing is likely to replace array-based genotyping in the future. To test the feasibility of schemes C and D (i.e. genotype the cows by SWGS rather than with SNP arrays, without (C) and with (D) imputation), we collected samples from a farm with 120 Holstein-Friesian cows. All cows were genotyped with the Illumina LD array (17K) as well as sequenced at average 1.08 −fold depth (range: 0.26-1.73). The milk was sequenced at ~3.5-fold depth. The correlation between predicted and measured SCC was 0.97 (or 0.96 when ignoring one cow with SCC > 3 million) under scheme A. Under scheme C, correlations were 0.82 (0.83) when sequencing the milk at 3.5x and 0.75 (0.76) when down-sampling the milk to 0.1x. We then imputed the sequenced cows to HD (770K SNPs) using a population of 800 reference animals genotyped with the HD array (scheme D). The correlation increased to 0.93 (0.94) when sequencing the milk at 3.5x and to 0.83 (0.77) when down-sampling the milk to 0.1x (Fig. 3C). Accuracies at SCC threshold of 500,000 were 0.96 (scheme A), 0.95 (3.5x) and 0.80 (0.1x) (scheme B), 0.82 (3.5x) and 0.81 (0.1x) (scheme C), and 0.95 (3.5x) and 0.88 (0.1x) (scheme D) (Fig. 3C). In summary, (i) combining cow genotyping using SNP arrays with genome-wide imputation with SWGS of tank milk allows for cost-effective identification of cows with subclinical mastitis even in farms with as many as 500 cows per milk tank, and (ii) as sequencing costs continue to decline, arrays-based targeted SNP genotyping of the cows could be replaced by genotyping by SWGS and yield comparable results.

### Monitoring SCC dynamics with the proposed method

Farmers typically measure individual SCC once a month or less. Yet, SCC may rapidly change. The SCC measured on the milk testing date may not be a reliable indicator of the cow’s udder health during the intervening period. To examine the SCC dynamics over time, we collected 20 tank milk samples over a 100-day period (day −84 to +17 from day of milk testing) for the farm with 120 cows. Milk samples were genotyped using the Illumina LD array, and individual SCC estimated using scheme A. Fig. 4A shows the SCC predicted every 5 days on average for the 120 cows, sorted by SCC measured on day 0 (=milk testing day). Of note, the correlation between the SCC measured on day 0 and the average of the SCC estimates for the 21 collection dates was low (*r* = 0.52)(Fig. 4B) and decreased rapidly with the number of days from milk testing day (Fig. 4C).

**Figure 4:**
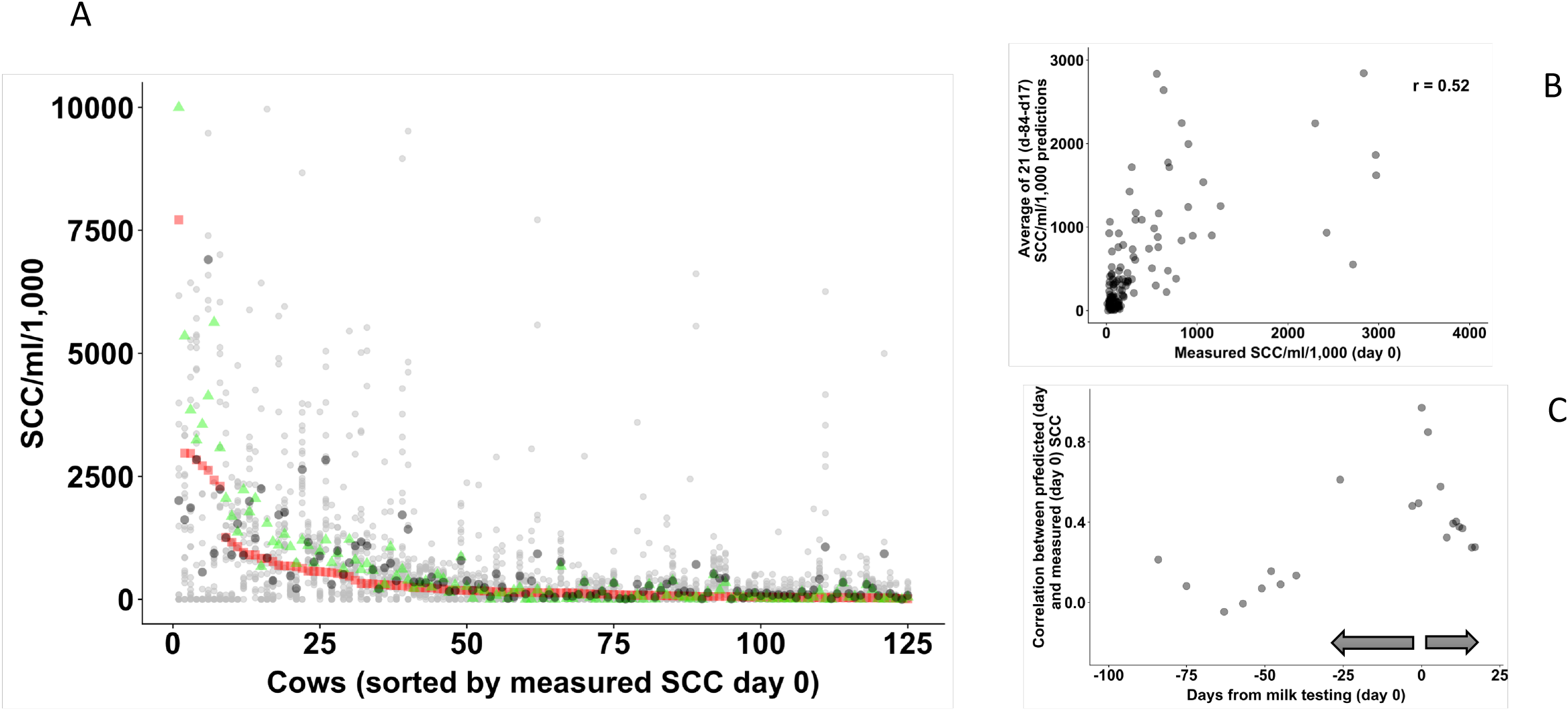
**(A)** SCC predicted using scheme A for 21 tank milk samples collected over a 100-day period from 125 cows total. Small grey circles: 20 predictions per cow. Large grey circles: average of 21 measurements per cow. Red square: SCC measured on day 0. Green triangle: SC<u>C</u> predictions on day 0. **(B)** Relationship between SCC values measured on day 0 and average of 21 predictions sampled over a 100-day period (days −84 to +17). **(C)** Correlations between measured (day 0) and predicted (day x) SCC as a function of the number of days from day 0.

## Discussion

We herein demonstrate that by combining array-based SNP genotyping and whole-genome imputation for the cows with SWGS of the tank milk, it is possible to accurately estimate SCC for individual cows and hence effectively identify animals with subclinical mastitis even for tanks collecting milk for >500 cows, and this by performing a single analysis for the entire herd. Reagent costs to sequence a mammalian genome at 1-fold depth are now <20€ thus making this a cost-effective proposition. As a matter of fact, the method is being deployed in the field in several countries.

Implementing the method requires all cows on the farm to be genotyped. This will increasingly correspond to reality as genotyping costs continue to decrease and genomic selection is more and more used for the selection of cows. In 2016 more than 1.2 million dairy cows had been reportedly genotyped in the US alone^8^ and present worldwide numbers are likely ≥ 3 million. In addition, a reference population of a few hundred animals of the breed of interest that are either HD genotyped (700K) or better whole-genome sequenced are required for accurate imputation. Such reference populations are already available for the most important dairy cattle breeds^7,9^, and could be easily generated for the remaining ones.

We show that SCC are dynamic and rapidly change over time. SCC measured on day 0 are poor indicators of SCC in previous and future weeks: cows with high SCC on the day of milk testing may have low S<u>C</u>C a few days later (or earlier) and vice versa. The proposed method would allow tighter monitoring of SCC hence improving udder health management. More frequent monitoring of SCC for large number of cows may reveal interindividual differences with regards to SCC dynamics that may be correlated with mastitis resistance, heritable and hence amenable to selection including by GS. Sequencing of the DNA in the tank milk allows simultaneous characterization of the tank’s microbiome. As a matter of fact, ~1% of reads in this study mapped to bacterial genomes (data not shown). This information may be very useful both from a farm health management point of view as well as from a downstream dairy processing point of view. Whole genome sequence data of bulk milk also informs about the herd frequency of functional variants such casein variants affecting consumer health or processing properties^10^, or variants causing inherited defects or embryonic lethality in cows^4^. In many countries, it is not allowed to add milk from cows being treated with antibiotics to the tank. As suggested before, the proposed approach can be adapted to verify whether a specific cow did contribute milk to the tank or not (f.i. by testing the significance of the corresponding cow effect in the linear model) ^3^. The described method may have applications in tracing the origins of bulk animal food products other than milk, as well as in monitoring the composition of mixed-donor blood-derived transfusion products.

## Acknowledgements

This work was funded by the Unit of Animal Genomics and by the ERC DAMONA grant to Michel Georges. We are grateful to Jean-Bernard Davière, Pierre Lenormand, Bonny Van Ranst, Kristien Neyens and Miel Hostens for providing the samples and information needed to conduct the experiments. <u>The proposed method is the subject of awarded (WO/2013/079289) and filed patents (PCT/EP2019/057628)</u>.

## Methods

### Simulated data

<u>Reference scheme (A)</u>: We simulated farms with *n* (25, 50, 100, 250 and 500) cows contributing milk to the tank. Cows were genotyped with SNP arrays for *m* (10K, 50K, or 750K) markers without error. Minor Allele Frequencies (MAFs) were sampled from a uniform [0,0.5] distribution, and genotypes from the corresponding Hardy-Weinberg distributions. SCC of individual cows (*SCC_i_*) were simulated by sampling values from a Weibull distribution with scale parameter *α*=1 and shape parameter *β*=2, and multiplying the ensuing value by 200,000. Exact B-allele frequencies of individual SNPs (*BAF_j_*) in the milk were determined for each SNP *j* based on the combination of cellular contribution of the *n* cows to the milk, and their genotype. It was assumed that B-allele frequencies were estimated with a normally distributed error *N*(0,0.0025) (i.e. SE = 0.05), yielding *m* 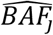. <u>Scheme B:</u> Same setting as in the reference scheme with the following additions. For cows genotyped for 10K or 50K SNPs, we simulated imputation by augmenting the data to 8 million (M) genotypes using an error model mimicking real, MAF-dependent imputation accuracy. The error model was constructed using a real data set for 800 unrelated Holstein-Friesian individuals that were genotyped for the Illumina 777K array. This data set was split into a set of 200 and a set of 600 individuals. The set of 200 was reduced first to the genotypes interrogated by the Illumina 10K (LD) array and then to the genotypes interrogated by the Illumina 50K SNP arrays. The reduced SNP sets were imputed back to the content of the Illumina 777K (HD) SNP array using the 600 individuals as reference population. The frequencies of imputing a given genotype depending on the real genotype, were scored for MAF bins of 0.01 separately for the LD and 50K array data. We simulated genotyping-by-sequencing of tank milk as follows. For each of the 8M SNP positions, we sampled local read depth (*r* ∈ integers) from a Poisson distribution with mean *C*, where *C* is the average genome-wide coverage (0.25, 0.5, 1, 2 or 5). We then sampled *r* reads, each with a probability = *BAF_j_* (computed as above) of being the B-allele. <u>Scheme C</u>: Individual SNP genotypes and tank B-allele frequencies (*BAF_j_*) were generated as in scheme A (genotypes at 8 M SNP positions). It was assumed that milk tank was genotyped by SWGS at average coverage of *C* (0.25, 0.5, 1, 2 or 5) and cows were genotyped by SWGS at average coverage of *C* (0.25, 0.5, or 1). Genotyping-by-sequencing of individual cows was simulated by (i) sampling, for each of 8M SNP positions, local read depth (*r* ∈ integers) from a Poisson distribution with mean *C*, and (ii) sampling *r* reads with probability 0, 0.5 or 1 to be the alternate allele (<u>B</u>) depending on the genotype of the cow (<u>AA</u>, <u>AB</u> or <u>BB</u>). Genotyping-by-sequencing of the tank milk was done as in Scheme A. <u>Scheme D</u>: Identical to scheme C except that cow genotypes were generated at 8M SNP position using a MAF- and sequence-depth dependent imputation error model. The error model was constructed using available SWGS data down sampled to 1x (176 cows) or 0.25x coverage (192 cows). The cows were imputed to HD (777K SNPs) using a reference population of 800 unrelated Holstein-Friesian individuals that were genotyped with the Illumina 777K array. At each of the 777K SNP positions, the likelihood of the sequence data under the three possible genotypes (<u>AA</u>, <u>AB</u> and <u>BB</u>), were computed following Chan et al.^3^, as:

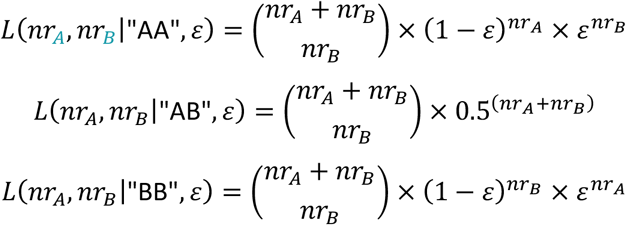

where *nr_A_* (respectively *nr*_B_) is the number of A (respectively B reads) and *ε* is the sequencing error rate set at 0.01. The corresponding log_10_ *L* were used as input for Beagle4^1^. Variant positions without sequence coverage in any of the 176 (192) cows (hence not imputed by Beagle4) were dealt with in a second round of imputation using Beagle5^2^. The imputation accuracy was evaluated in 0.01 MAF-bins by comparing imputed and real genotypes at the ~17K variant positions interrogated by the Illumina LD array.

### Real data

<u>Data set 1</u>: We obtained a sample of tank milk from a farm in France milking 133 Holstein-Friesian cows. All had been genotyped with an Illumina LD array interrogating 17K SNPs using standard procedures. For all cows, genotypes were imputed to whole genome using a reference population of 743 Holstein-Friesian animals sequenced at average depth of 15x (range: 4-48) and the Beagle software (v5.0)^1^ yielding allelic dosages for a total of 13 million SNPs. Individual milk records, including volume and SCC (cells/ml) measured on the day of the sample collection, were obtained for all cows that had contributed milk to the tank. DNA was isolated from 1.5 ml tank milk using the NucleoMag kit (Macherey-Nagel). The tank milk DNA was first genotyped using the Illumina LD array interrogating 17K SNPs. An Illumina compatible NGS library was then prepared with 50ng of genomic DNA using the KAPA HyperPlus kit (Roche). Sequencing was performed on a NextSeq500 instrument (Illumina), yielding 63 million paired end reads of 2*75 bp, corresponding to a genome coverage of 3.5x. Reads were mapped to the bosTau8 genome build using BWA mem. Reference (R) and alternate (A) alleles were counted at 13M SNP positions of the HD array using the Bam-ReadCount tool (https://github.com/genome/bam-readcount.git) for reads with a minimum mapping quality of 30. <u>Data set 2</u>: We obtained samples of tank milk from a Belgian farm including milk from 520 Holstein-Friesian cows. Milk volume and SCC (cells/ml) measured on the same day, were obtained for all cows that had contributed milk to the tank. All cows were genotyped with the Illumina LD array interrogating 17K SNPs using standard procedures, and imputed to whole genome using whole genome sequence data (average depth: 15x; range: 4x-48x) from 743 Holstein-Friesian animals as reference (M. Georges, unpublished) and the Beagle software (v5.0)^2^ yielding allelic dosages for a total of 13 million SNPs. DNA extraction from the tank milk samples and genotyping with the Illumina LD (17K) array were conducted as for dataset 1. For sequencing of the tank milk, an illumina compatible sequencing library was prepared using 12 ng of DNA and the Riptide High Throughput Rapid Library Prep Kit (iGenomx). The library was sequenced on an Illumina NextSeq500 2*150 paired end flow cell at 4X coverage. <u>Data set 3</u>: We obtained samples of tank milk from a Belgian farm including milk from 120 Holstein-Friesian cows. Milk volume and SCC (cells/ml) measured on the same day, were obtained for all cows that had contributed milk to the tank. All cows were genotyped with the Illumina LD array interrogating 17K SNPs using standard procedures, and imputed to whole genome using whole genome sequence data (average depth: 15x; range: 4x-48x) from 743 Holstein-Friesian animals as reference (M. Georges, unpublished) and the Beagle software (v5.0)^2^ yielding allelic dosages for a total of 13 million SNPs. We additionally prepared Illumina compatible NGS library for each cow, using 12 ng of genomic DNA and the Riptide High Throughput Rapid Library Prep Kit (iGenomx). Libraries were sequenced on an Illumina Novaseq S4 2*150 paired end flow cell at average 1.08x depth (range: 0.26x-1.73x). Cow genotype-by-sequencing data were imputed to HD (777K) density using a reference population of 800 Holstein-Friesian animals genotyped with the bovine HD Illumina array (777K SNPs) and the Beagle software (v5.0)^2^ yielding allelic dosages for a total of 777K SNPs. DNA extraction from the tank milk samples, genotyping with the Illumina LD (17K) array, and sequencing (coverage 4x) were conducted as for datasets 1&2. <u>Data set 4</u>: In addition to obtaining a sample of tank milk on the day of the milk recording (i.e. yielding the SCC measured using with a cell counter) for the Belgian farm with 120 cows, we weekly collected an additional 11 tank milk samples before and 9 samples after, spanning a total period of ~3 months. The corresponding DNA samples were genotyped using the Illumina LD (17K) array.

### Statistical model

We defined a set of *m* linear equations of the form:

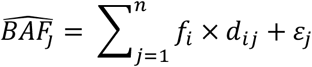

in which *f_i_* is the proportion of the DNA in the tank milk contributed by cow *i*, *d_ij_* is the “dosage” of the alternate allele A for cow *i* and marker *j*, and *ε_j_* is the error term for marker *j*. When genotyping the tank milk with arrays, 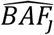 corresponds to the B-allele frequency estimated by Genome Studio (Illumina). When genotyping the tank milk by SWGS, 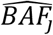 corresponds to the proportion of A reads at the corresponding genome position. For cow genotypes obtained with arrays, *d_ij_* corresponds to 0, 0.5 or 1 for genotypes AA, AB and BB, respectively. For cow genotypes obtained by imputation, *d_ij_* is the dosage of the B allele estimated by Beagle. For cow genotypes obtained by SWGS, 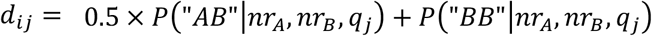 where *nr_A_* (respectively *nr*_B_) is the number of A (respectively B reads) for marker *j* and cow i, and *q_j_* is the population frequency of the B allele of marker j.

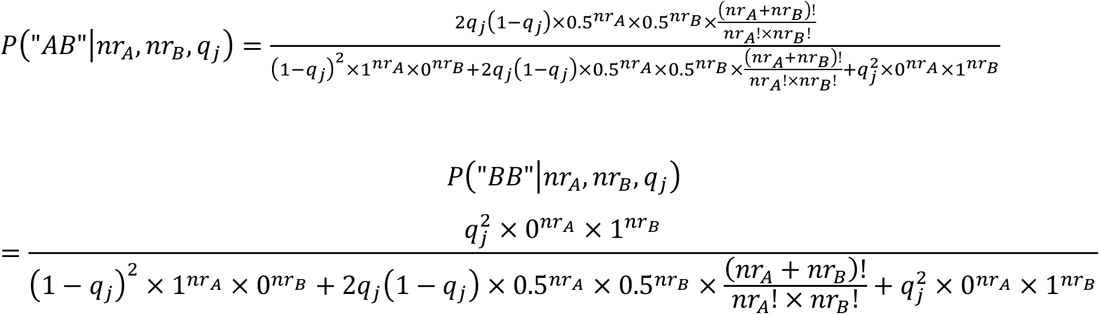

For SNPs *j* without usable information for cow *i* (f.i. genotyping failure or no covering reads) *d_ij_* was set at 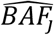.

The *f_i_’*s were estimated by least square analysis, i.e. by minimizing 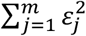. When the tank milk was genotyped by SWGS, we also performed a weighted least square analysis, i.e. we estimated *f_i_’*s by minimizing 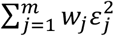, where *w_j_* is the coverage (*nr_A_* + *nr_B_*).

The *SCC_i_’*s were calculated from the *f_i_’*s

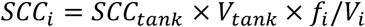

Where *V_tank_* and *V_i_*, are the volumes of milk in the tank and contributed by cow i, respectively.

The accuracies of the predictions were measured by the (i) correlation (*r*) between real and estimated *SCC_i_*, and/or (ii) the ability to discriminate animals with SCC above versus below a certain threshold value measured as (*T_P_* + *T_N_*)/*n*, where *T_P_* stands for the number of true positives, *T_N_* for the number of true negatives, and *n* for the total number of cows.

To test the effect of sequence depth on accuracy we sampled reads overlapping SNP positions with probability *x*, such that *E*(*C × x*) = *D*, where *D* is the desired sequence depth.

